# Instrumented immobilizing boot quantifies reduced Achilles tendon loading during gait

**DOI:** 10.1101/2020.02.27.968495

**Authors:** Todd J Hullfish, Kathryn M. O’Connor, Josh R. Baxter

**Affiliations:** Department of Orthopaedic Surgery, University of Pennsylvania, Philadelphia, Pennsylvania, USA

**Keywords:** plantar flexor, biomechanics, insole, rupture, gait, personalized medicine

## Abstract

Achilles tendon ruptures are common injuries that lead to functional deficits in two-thirds of patients. Progressively loading the healing tendon has been associated with superior outcomes, but the loading profiles that patients experience throughout rehabilitation have not yet been established. In this study, we developed and calibrated an instrumented immobilizing boot paradigm that is aimed at longitudinally quantifying patient loading biomechanics to develop personalized rehabilitation protocols. We used a 3-part instrumented insole to quantify the ankle loads generated by the Achilles tendon and secured a load cell in-line with the posterior strut of the immobilizing boot to quantify boot loading. We then collected gait data from five healthy young adults to demonstrate the validity of this instrumented immobilizing boot paradigm to assess Achilles tendon loading during ambulation. We developed a simple calibration procedure to improve the measurement fidelity of the instrumented insole needed to quantify Achilles tendon loading while ambulating with an immobilizing boot. By assessing Achilles tendon loading with the ankle constrained to 0 degrees and 30 degrees plantar flexion, we confirmed that walking with the foot supported in plantar flexion decreased Achilles tendon loading by 60% (*P* < 0.001). This instrumented immobilizing boot paradigm leverages commercially available sensors and logs data using a small microcontroller secured to the boot and a handheld device, making our paradigm capable of continuously monitoring biomechanical loading outside of the lab or clinic.

## INTRODUCTION

Over the past 3 decades, acute Achilles tendon ruptures have increased 10-fold (Lantto et al., 2015), leading to plantar flexion deficits in two out of three patients one year after injury (Brorsson et al., 2017). Tendon elongation and plantar flexor muscle remodeling explain these poor functional outcomes in patients (Hullfish et al., 2019; Silbernagel et al., 2012) and are likely caused by suboptimal loading throughout healing (Hillin et al., 2019; Williams, 1990). To protect the healing tendon from excessive loading, patients wear immobilizing boots for 8-12 weeks following the initial treatment of the Achilles tendon rupture, whether treated surgically or nonsurgically (Willits et al., 2010). However, the mechanical loads applied to the Achilles tendon during ambulation in these immobilizing boots are unknown, limiting the potential efficacy of rehabilitation protocols.

To address this unmet clinical need, we developed an instrumented immobilizing boot paradigm to quantify Achilles tendon loading and boot load sharing during ambulation. Using an immobilizing boot currently prescribed in our clinical practice, we secured a 3-part load sensitive insole inside the boot to quantify loads transfered through the foot by the Achilles tendon and instrumented the posterior strut with a load cell to quantify boot loading. In this technical note, we describe the technical aspects we considered as we developed this instrumented immobilizing boot paradigm and present experiments establishing its validity.

## METHODS

### Instrumented immobilizing boot

We developed an instrumentation paradigm to quantify Achilles tendon loading during ambulation in an immobilizing boot used to treat Achilles tendon ruptures (**Figure 1**). To do this, we decided to leverage a commercially available immobilizing boot (Vacoped, Oped, Munich, Germany) that is currently prescribed in our clinics. Our framework relies on two assumptions: 1) that internal plantar loading measured by the instrumented insole is generated by the Achilles tendon acting to plantar flex the foot; and 2) that the rigid strut carries all the reaction load between the boot and the ground. In addition to being used in our clinics, this immobilizing boot has several other characteristics that improved our instrumentation paradigm: 1) ankle motion is constrained by a posteriorly-positioned rigid plastic strut that is kinematically deterministic, which allows us to calculate the moment carried by the boot and 2) the boot is made of plastic, which allows for additional electronics to be rigidly secured to the boot.

**Figure 1.**
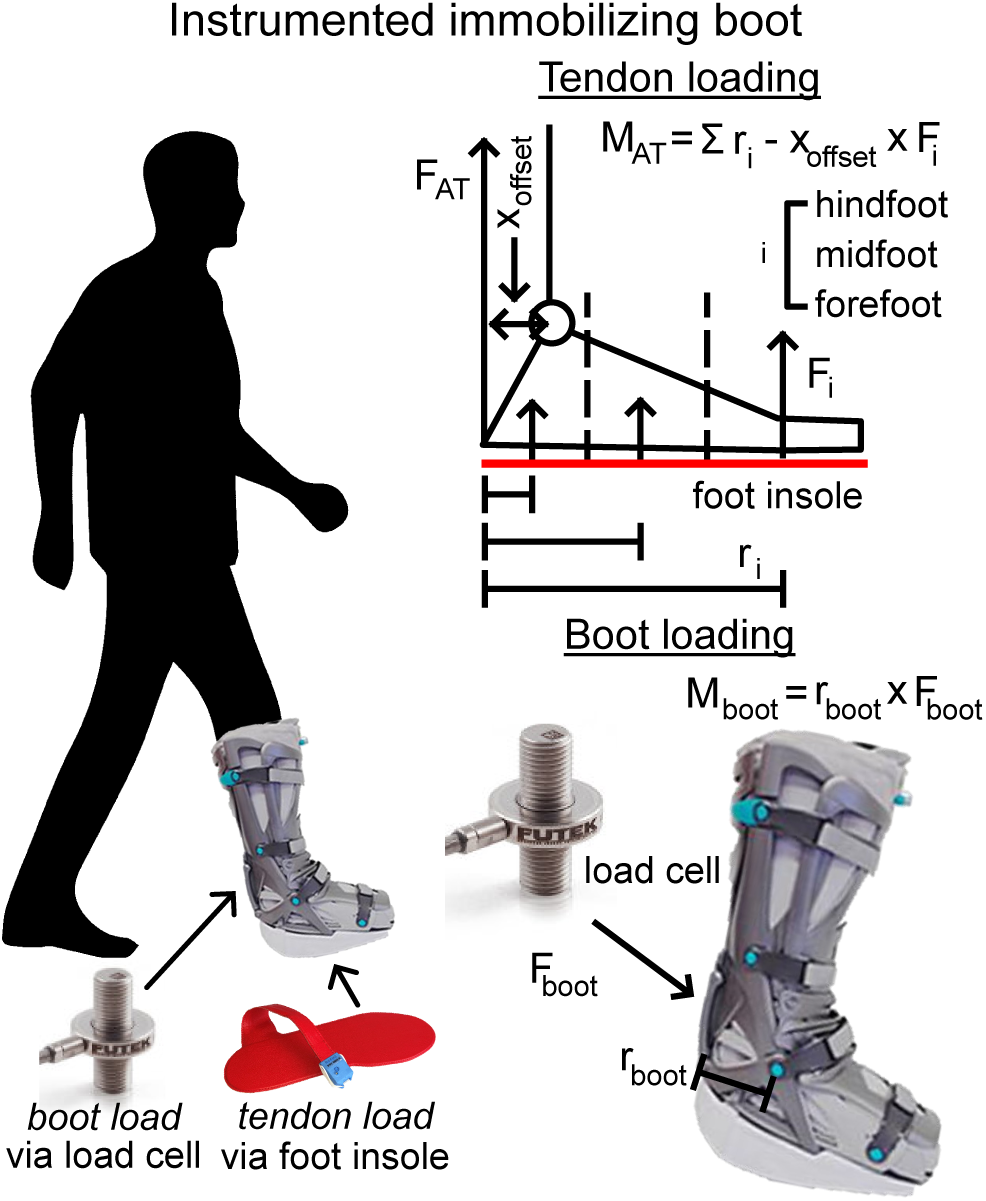
Our instrumented immobilizing boot paradigm quantified internal loads carried by the Achilles tendon and external loads carried by the boot. We used an instrumented insole (top) to quantify the sagittal loading of the ankle joint, which can be transformed to Achilles tendon load. The loads carried by the boot were calculated as the product of the moment arm and posterior strut force, which we measured using an inline loadcell. This net boot load represented the sum of the internally generated Achilles tendon load and the externally applied ground reaction force during gait.

To quantify Achilles tendon loading during ambulation, we placed a 3-part force-sensitive shoe insole (Loadsol, Novel Electronics, St. Paul, MN) under the vacuum boot liner. In our recent work, we demonstrated that the instrumented insole accurately quantifies plantar flexor loading over a wide range of activities including walking, running, and jumping in normal running shoes (Hullfish and Baxter, 2019). In this study, we extended our implementation of this instrumented insole into an immobilizing boot to quantify Achilles tendon loading during walking. To ensure that all loads measured by the instrumented insole were the result of active plantar flexion, we tared the insole with the boot unloaded from the ground. Because the boot itself is passive, any loads measured by the insole result from internal work done by the plantar flexors.

To quantify boot loading during ambulation, we measured the load carried by the posterior strut using a load cell (LCM200, Futek, Irvine, CA) secured in-line with the strut (**Figure 1**). We calibrated the load cell with known loads on a mechanical testing frame and confirmed that it had a strongly linear response to loading (R2 > 0.99). By calculating the product of the boot lever arm and the posterior strut load cell, we determined the net loads carried by the boot. This net load was the summation of the load carried by the strut that resisted the internally generated loads from the Achilles tendon and the externally applied ground reaction force during gait. We conditioned these strain gauge measurements using an amplifier (HX711, Sparkfun, Niwot, Colorado).

These measurements were logged using two discrete systems and synchronized to a common clock. Instrumented insole data was logged to a hand-held device (iPod Touch, A1574, Apple, Cupertino, CA) using an app provided by the insole vendor. The posterior strut load cell was logged to a memory card on the microcontroller (FeatherSD, Adafruit, New York, NY) that controlled the amplifier. We synchronized both systems by having subjects cyclically load and unload the immobilizing boot on top of a force plate 5 times. Using the 5 loading peaks, we calculated time offsets between these three systems and determined a time offset that we applied to our system master clock.

### Gait analysis and validation

Five healthy young adults (3 males, 2 females, 30 ± 5.5 years, 29.4 ± 8.1 BMI) participated in this IRB approved study and provided written informed consent. Participants walked at a self-selected speed while wearing the immobilizing boot at 2 conditions: 1) ankle position set to 30 degrees plantar flexion and 2) ankle position set to 0 degrees neutral. We also collected motion capture and ground reaction force data during these test conditions. In this study, we secured markers to the medial and lateral aspects of the immobilizing boot to track the positions of the boot that represented the ankle joint axis, toe joint axis, tibial mid-shaft, and posterior strut. We calculated total boot moment as the cross product between the ground reaction force and the boot joint that was in line with the biologic ankle joint. During these trials, subjects wore a rubber shoe lift (EvenUp, Oped, Munich, Germany) on the contralateral foot to compensate for the thickness of the boot bottom.

We developed an algorithm to calibrate the instrumented insole based on external and internal loads we measured with motion capture, the instrumented boot, and the instrumented insole. We assumed that the moment measured by the instrumented immobilizing boot was equivalent to the difference of the externally applied load from the ground reaction force and the internally applied plantar flexor moment (Eq. 1). We used the experimental measurements acquired during walking at 0 degrees plantar flexion to calibrate the instrumented insole for both test conditions. By directly measuring this net boot loading as the product of the boot load cell and the load cell moment arm (Eq. 2), we developed an objective function to minimize the measurement error of the Achilles tendon load measured by the instrumented insole (Eq. 3). Briefly, we used a gradient based solver (fsolve, MATLAB 2019b, The Mathworks Inc, Natick, MA) to minimize the root mean square error (RMSE) of the objective function by iteratively adjusting the geometric parameters of the instrumented insole (**Figure 1**). We provided initial guesses of the centers of pressure of the three instrumented insole sensors and the position of the ankle joint with respect to the posterior aspect of the insole. We hypothesized that this calibration algorithm would reduce Achilles tendon loading errors to below 10% of the estimated Achilles tendon load. We then used this calibrated instrumented insole to confirm that prescribing the ankle at 30 degrees plantar flexion (condition 1) offloads the Achilles tendon more than prescribing 0 degrees plantar flexion (condition 2).

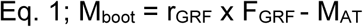

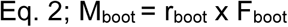

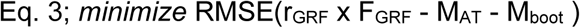

We calculated the boot load cell moment arm (r_boot_) as the distance from the load cell to the boot ankle hinge. Similarly, we calculated the ground reaction force moment arm (r_GRF_) as the distance from the ground reaction force center of pressure to the boot ankle joint, which we defined as the midpoint between the lateral and medial ankle markers. We calculated the moment generated by the Achilles tendon (M_AT_) using our validated instrumented insole algorithm (Hullfish and Baxter, 2019).

## RESULTS

During walking at a self-selected speed with the immobilizing boot constrained to 0 degrees plantar flexion, subjects consistently loaded their Achilles tendons (**Figure 2**). By iteratively adjusting the geometric parameters of the instrumented insole, we decreased our objective function by 78% (uncalibrated RMSE 13.6%, calibrated RMSE 3.0%, **Figure 2 bottom**). This calibration decreased expected instrumented insole errors nearly approximately 16%. After applying these calibration factors, we detected a 60% decrease in Achilles tendon loading when the foot was placed in 30 degrees plantar flexion compared to 0 degrees plantar flexion (*P* < 0.001, **Figure 3**).

**Figure 2.**
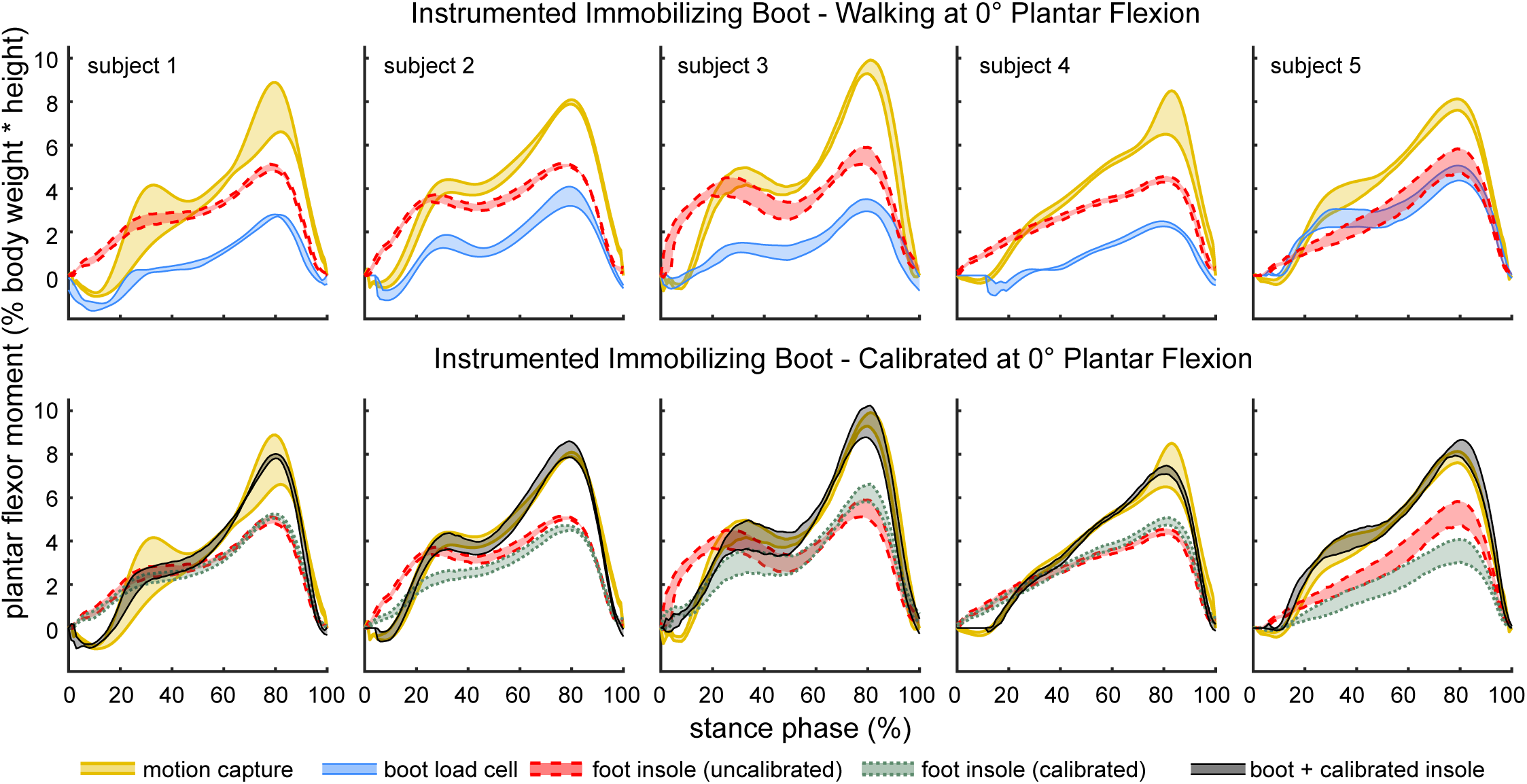
We measured and normalized moments using motion capture (thick gold), the immobilizing boot load cell (thin blue), and the instrumented insole (dashed red) (top row). To calibrate the instrumented insole, we assumed that the moment measured directly by the immobilize boot load cell was equivalent to the difference between the externally applied moment and instrumented insole. We used a gradient based optimization routine (fsolve, MATLAB 2019a) to iteratively adjust the geometric parameters of the instrumented insole. This optimization approach converged on solutions (bottom row) and had noticeable effects on the plantar flexor moment approximated from the foot insole (dashed green). We visually confirmed that the optimization routine converged by plotting the normalized motion capture moment (thick gold) and the sum of the moments measured by the instrumented boot and calibrated insole (thin black). Shaded areas are 95% boot strapped confidence intervals.

**Figure 3.**
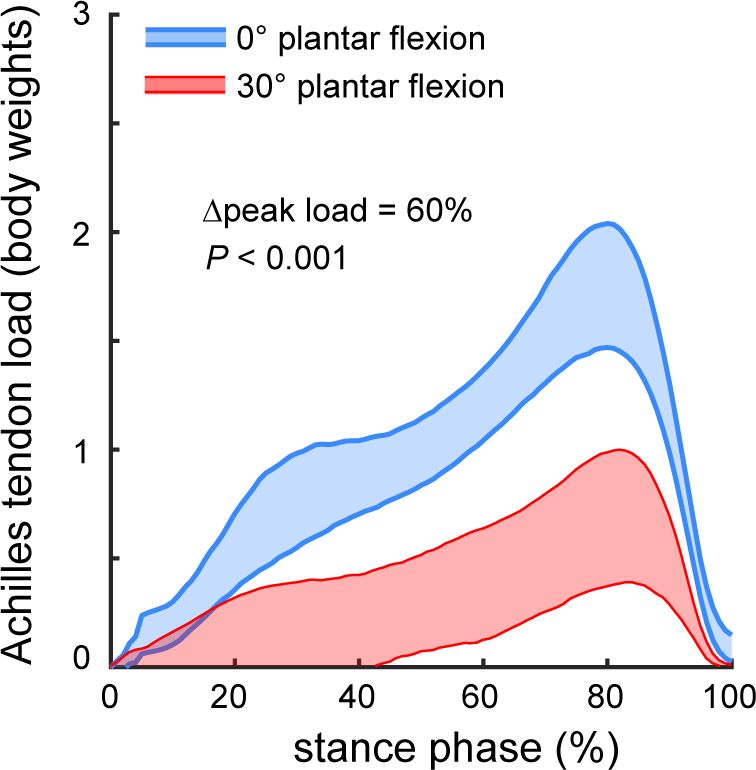
By wearing the immobilizing boot constrained at 0 degrees plantar flexion (thin red), peak Achilles tendon loads were reduced by 60% compared to walking with the ankle constrained at 30 degrees plantar flexion (thick blue). Shaded areas are 95% boot strapped confidence intervals.

## DISCUSSION

In this study, we developed an instrumented immobilizing boot paradigm that addresses an unmet clinical need. By using a commercially available instrumented insole and load cell, we can now quantify loads carried by both the biologic tissue and immobilizing boot during ambulation. Further, this paradigm logs data continuously, which permits patient loading data to be captured away from the clinic or research lab. Monitoring patient loading provides clinicians and researchers critical new data to understand the mechanical mechanisms that govern muscle remodeling and tendon healing following Achilles tendon rupture. This approach will also add quantitative data to accelerated rehabilitation paradigms, which is currently limited by a disconnect between actual patient loading and perceived patient loading (Aufwerber et al., 2019).

Our paradigm demonstrates the large amount of Achilles tendon loading generated during gait in an immobilizing boot, which is not biomechanically necessary because the boot position is prescribed by the rigid plastic strut. Compared to shod walking at a comfortable speed, walking in an immobilizing boot at 0 degrees plantar flexion only reduces Achilles tendon loading by approximately 50% (Hullfish and Baxter, 2019). Earlier work found that increasing the amount of ankle support in a heel-wedge boot decreases the amount of plantar flexor muscle activity (Zellers et al., 2019). As patients progress along their rehabilitation, they load their Achilles tendon with greater frequency and magnitude (Aufwerber et al., 2019). However, no studies to date have precisely quantified Achilles tendon loading throughout healing. Our instrumented immobilizing boot paradigm provides researchers with a new framework linking clinical outcomes with longitudinal tendon loading. This instrumented immobilizing boot paradigm has several limitations that need to be addressed. Our current paradigm requires external loads measured during walking to calibrate the instrumented insole. We are currently developing a simplified calibration procedure that does not require a motion capture system. We assumed that all loads measured by the instrumented insole were generated by the Achilles tendon force. However, future research is needed to confirm this assumption in patients with Achilles tendon ruptures who exhibit different neuromuscular control (Suydam et al., 2015). Our future work will longitudinally monitor patients healing from tendon ruptures to determine the loading profiles that govern muscle-tendon healing and functional outcomes.

In conclusion, we developed and validated an instrumented immobilizing boot paradigm that quantifies the loading biomechanics of both the Achilles tendon and immobilizing boot. This paradigm can also be extended to monitor tendon loading in patients with Achilles tendinopathy who wear ankle-foot orthoses. By continuously monitoring these loading profiles, researchers and clinicians will be able to identify how rehabilitation loading governs muscle-tendon healing and functional outcomes. This is a necessary step towards personalized medicine approaches for patients with tendon pathologies.

## ACKNOWLEDGEMENTS

We thank Oped Medical, Inc for providing the immobilizing boot used in this study.

